# MinD2 modulates cell shape and motility in the archaeon *Haloferax volcanii*

**DOI:** 10.1101/2024.08.01.606218

**Authors:** Megha Patro, Shamphavi Sivabalasarma, Sabrina Gfrerer, Marta Rodriguez-Franco, Phillip Nußbaum, Solenne Ithurbide, Sonja-Verena Albers

## Abstract

In bacteria and archaea, proteins of the ParA/MinD family of ATPases regulate the spatiotemporal organization of various cellular cargoes, including cell division proteins, motility structures, chemotaxis systems, and chromosomes. In bacteria, such as *Escherichia coli*, MinD proteins are crucial for the correct placement of the Z-ring at mid-cell during cell division. However, previous studies have shown that none of the 4 MinD homologs present in the archaeon *Haloferax volcanii* have a role in cell division, suggesting that these proteins regulate different cellular processes in haloarchaea. Here, we show that while deletion of MinD2 in *H. volcanii* (Δ*minD2*) does not affect cell growth or division, it impacts cell shape and motility by mispositioning the chemotaxis arrays and archaellum motors. Finally, we explore the links between MinD2 and MinD4, which has been previously shown to modulate the localization of chemosensory arrays and archaella in *H. volcanii*, finding that the two MinD homologues have synergistic effects in regulating the positioning of the motility machinery. Collectively, our findings identify MinD2 as an important link between cell shape and motility in *H. volcanii* and further our understanding of the mechanisms by which multiple MinD proteins regulate cellular functions in haloarchaea.

## Introduction

A proper spatial distribution of cellular components is essential for the optimal functioning of cells. In bacteria and archaea, the ParA/MinD family of ATPases is crucial for the spatiotemporal organisation of various cellular cargoes. For example, ParA proteins are involved in plasmid partitioning and chromosome segregation (Baxter and Funnell, 2014; Jalal and Le, 2020), while MinD is known for its role in regulating the placement of the bacterial divisome (Lutkenhaus, 2007). However, ParA/MinD proteins are not restricted to these functions and have been shown to modulate the positioning of several other cellular components, including flagella (Pulianmackal et al., 2023), chemotaxis systems (Ringgaard et al., 2011), and the conjugation machinery (Atmakuri et al., 2007).

Although several ParA/MinD homologs are encoded in archaeal genomes, their distribution and functions are still being elucidated. Notably, while ParA can be found in almost all archaea, only Euryarchaeota encode for MinD homologs (Nußbaum et al., 2020). Furthermore, although a few structural analyses have been carried on archaeal MinD homologs (*Pyrococcus horikoshii, Pyrococcs furiosus* and *Archaeoglobus fulgidus*) (Jeoung et al., 2009; Szklarczyk et al., 2017), the functional roles of MinD have so far only been studied in *H. volcanii* (Nußbaum et al., 2020). *H. volcanii* encodes for 4 MinD homologs: *minD1* (HVO_0225), *minD2* (HVO_0595), *minD3* (HVO_1634) and *minD4* (HVO_0322). Notably, in contrast to the critical role of MinDs in regulating cell division in bacteria, deletion of all 4 MinD homologs had no role in cell division or growth in *H. volcanii* (Nußbaum et al., 2020). However, deletion of MinD4 reduced archaeal swimming motility due to the mispositioning of chemotaxis arrays and archaellum motors, suggesting a significant role of MinD4 in governing these processes (Nußbaum et al., 2020). However, the role of other MinD homologues remains unclear.

In this study, we characterised the functions of the MinD2 protein of *H. volcanii*. Since MinD2 was observed to not directly affect FtsZ localisation (and thus cell division) or cell growth, it suggests a role in other cellular pathways (Nußbaum et al., 2020). Using genetic mutants, we showed it has a crucial role in determining cell shape, influencing the transition from rod-shaped to plate-shaped cells. Furthermore, using fluorescently tagged MinD2 variants, we demonstrated that the protein has a diffused localisation pattern, which suggests a potential regulatory mechanism for its cellular functions, possibly involving interactions with partner proteins. Additionally, we demonstrated that MinD2 synergizes with MinD4 to modulate chemosensory array localisation, archaella assembly and motility, further illustrating the role of MinD2 homologues in spatial organisation. Overall, our findings contribute to a deeper understanding of the function of MinD2 in *H. volcanii*, highlighting its multifaceted role in coordinating cellular morphology and motility.

## Results

### MinD2 impacts cell shape

To elucidate the function of MinD2 in *H. volcanii*, we started by characterizing the growth of a mutant strain lacking MinD2 (Δ*minD2)*. Growth rate measurements showed that the Δ*minD2* strain exhibited a growth pattern similar to that of the wild-type (WT) strain H26 (Figure S1), suggesting that MinD2 does not directly influence growth in *H. volcanii* as previously shown by (Nußbaum et al., 2020).

Then, we assessed the impact of MinD2 on morphology, using phase contrast microscopy to compare the cell shape and size of WT vs Δ*minD2* cells during different growth stages. As previously described, we found that WT *H. volcanii* (H26) undergoes growth-dependent alterations in its cell morphology. The time point of this shape transition has been observed to differ greatly based on medium components and is also affected by the presence of plasmids (Li et al., 2019; de Silva et al., 2021; Patro et al., 2023). During the early log phase, the liquid culture is predominantly composed of rod-shaped cells (R), which gradually transition into an intermediate state (I) and then into plate-shaped cells (P) as the optical density (OD^600^) of the culture increases. The categorisation of cell shape was conducted as previously described in (Patro et al., 2023).Indeed, H26 (WT) cells predominantly exhibited rod-shaped morphology until an OD^600^ of 0.03, with a minority showing I or P shapes (R = 41%; I = 26%; P = 33%) (Figure 1 and S2a). Starting from an OD^600^ of 0.06, a noticeable transition towards P cells became evident, with plates representing the majority (P = 55%). This trend persisted as the culture progressed, with the frequency of P cells reaching 81% at an OD^600^ of 0.2. By contrast, the majority of Δ*minD2* cells exhibited a plate-shaped morphology even in the early log phase (OD^600^ 0.01), with few I and R cells (R = 17%; I = 33%; P = 50%) (Figure 1 and S2a). The frequency of P cells continued to increase at higher culture densities, with more than 80% of the cell population adopting a plate-shaped morphology at OD^600^ above 0.06 (Figure 1 and S2a).

**Figure 1.**
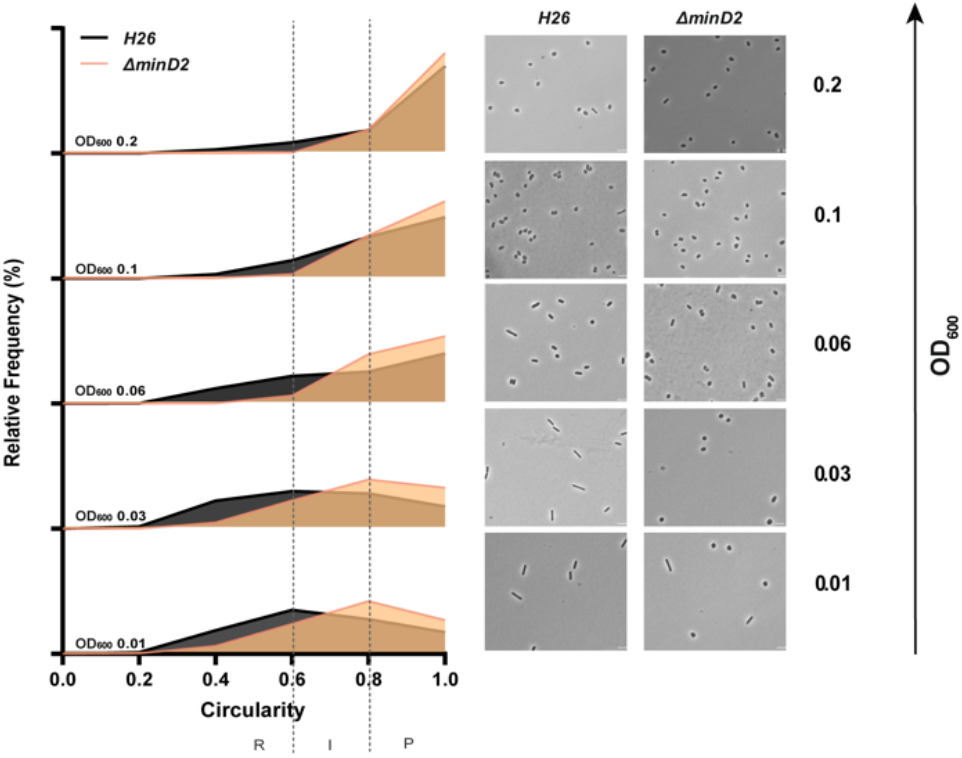
Cell shape analysis in Δ*minD2* strain and wild-type H26 cells throughout the growth curve. Left: Relative frequency distribution of cell circularity comparing H26 (grey) and Δ*minD2* (orange) analysed from micrographs. Vertical dashed line represents the different cell type R: Rods, I: Intermediates and P: Plates. Sum of the graph height per OD_600_ equals 100% and Y-axis indicates the percentage of cell population per cell type. Right: Phase contrast micrographs showing H26 and Δ*minD2* at different growth stages from OD_600_ 0.01 to 0.2 (bottom to top). Scale bar: 4μm. n_H26_ >1745, n_Δ*minD2*_ >1535. Three independent experiments with biological triplicates were carried out for both the strains.

Additionally, a comparison of the cell area in both strains displayed a similar trend, with Δ*minD2* cells having significantly smaller areas than WT cells at an OD^600^ of 0.01 (Figure S2b). Nevertheless, the areas of both WT and Δ*minD2* cells were comparable at higher densities (OD^600^ ≥ 0.03).

Collectively, these analyses demonstrate that while MinD2 does not affect cell growth, it significantly influences cell shape, with the deletion of *minD2* resulting in the loss of the ability to maintain a rod-shaped morphology, particularly in the early growth phase.

Previously, we found that the presence of a plasmid played a transient role in maintaining rod-shaped morphology in *H. volcanii* (Patro et al., 2023). To test this, we further examined the impact of MinD2 deletion on cell shape and size in the presence or absence of the empty plasmid pTA1392. In WT cells, the transition from rods to plates occurred between OD^600^ 0.06 to 0.1 when pTA1392 was present, as opposed to the transition occurring between OD^600^ 0.03 to 0.06 in the absence of pTA1392 (Figure S1b; Patro et al., 2023). Similarly, Δ*minD2* cells exhibited a higher population of rod-shaped cells at OD^600^ 0.01 in the presence of pTA1392 compared to Δ*minD2* without the plasmid. In Δ*minD2* + pTA1392, cells transitioned into intermediate (I) shapes (39%) at OD^600^ 0.03 and predominantly adopted a plate-shaped morphology (40%) at OD^600^ 0.06. Notably, this transition to a predominantly plate-shaped morphology occurred one generation time earlier in Δ*minD2* + pTA1392 (between OD^600^ 0.03 to 0.06) than in WT + pTA1392 (between OD^600^ 0.06 and 0.1). These findings further highlight the role of MinD2 in determining cell shape, particularly in promoting the ability of cells to maintain a plate-shaped morphology. As reported earlier (Patro et al., 2023), the presence of the plasmid increases the frequency of rod cells in the *minD2* mutant, suggesting that the plasmid induces rod development. Furthermore, while cells that contained plasmids maintained rod-shaped morphology longer throughout the different growth phases, the absence of *minD2* still reduced the ability to maintain a rod-shaped morphology throughout all phases (Figure S1b, S2c,d).

### MinD2 deletion impacts the assembly of archaella

In *H. volcanii*, cell shape is linked to swimming motility. During the early logarithmic phase, rod-shaped *H. volcanii* cells exhibit motility, powered by the assembly of polar bundles of archaella (Li et al., 2019). However, as cells transition into plate shape during stationary phase, the archaellum filaments are lost, resulting in motility cessation. Given the observed impact of MinD2 deletion on cell shape, we investigated whether the Δ*minD2* also showed changes in archaella assembly and motility. For this, we used transmission electron microscopy (TEM) to characterize WT and Δ*minD2* cells collected at an early log phase (OD^600^ 0.03). However, *H. volcanii* cells express several pili throughout different growth phases (Esquivel et al., 2016), which are difficult to differentiate in diameter and size from archaella. Therefore, we further deleted *pilB3*, the pilus assembly ATPase, in both the H26 and Δ*minD2* strain. These Δ*pilB3* mutants lack pili, which facilitates visualisation of archaella in these strains.

The results revealed that the rod-shaped H26Δ*pilB3* cells, both with and without plasmid pTA1392, displayed archaellation in 80% and 83.7% of the cells, respectively (Figure 2a and 2c). By contrast, the Δ*minD2*Δ*pilB3* strain, characterised by a discoid-shaped phenotype, only the few rod-shaped cells observed showed archaellation (∼20% of the cells). In the presence of pTA1392, where the percentage of rod-shaped cells is slightly increased (24%; Figure S2a and S2b), the percentage of archaellated cells is also observed to increase to ∼32% (Figure 2b and 2c).

**Figure 2.**
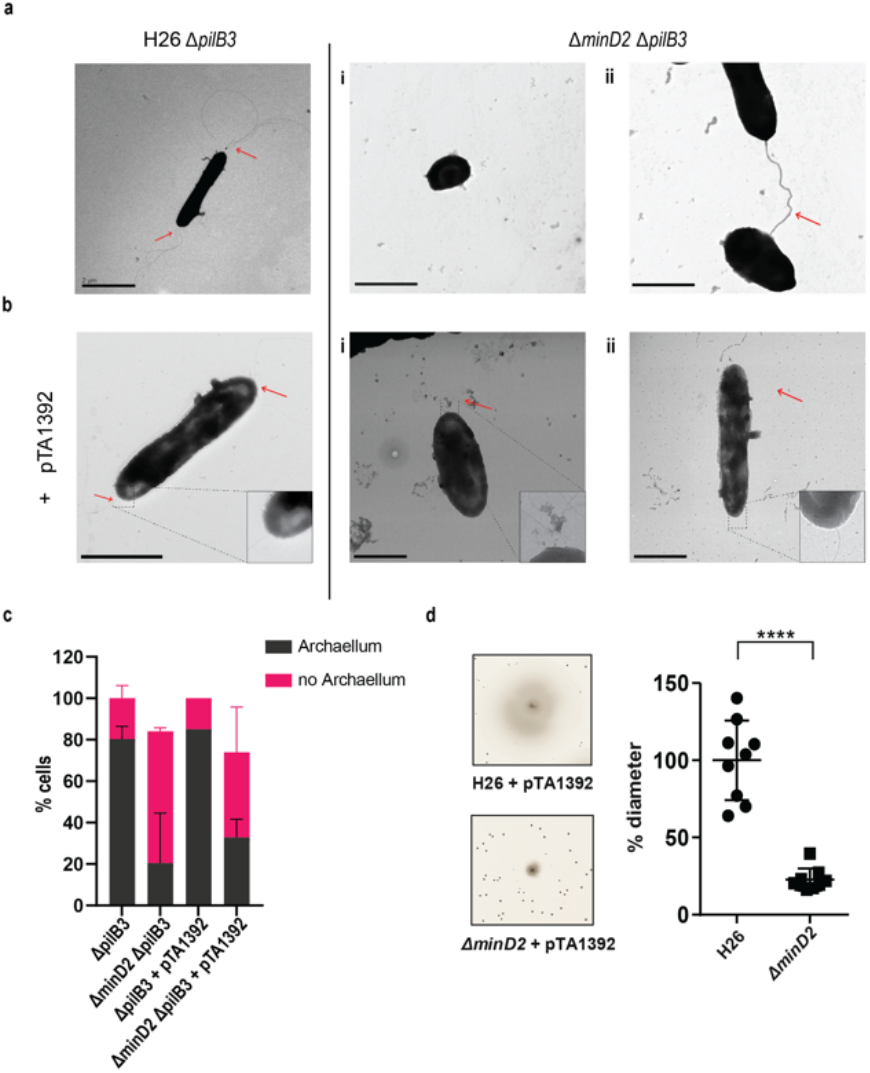
The deletion of *minD*2 affects cell motility and archaellation. (a) Transmission electron microscopy of H26 *ΔpilB3* cells showing archaella and *ΔpilB3ΔminD2* (i) plate-shaped cell without archaella and (ii) rod shaped cell with archaella (b) H26 *ΔpilB3* + pTA1392 cells showing archaella and *ΔpilB3ΔminD2* + pTA1392 (i) cell in transition rod to round (ii) rod shaped cell showing archaella. All cells were visualised at an early exponential phase (OD_600_: 0.03). Scale bar: 2µm (c) Distribution of cells with or without archaella for H26 ΔpilB3 and ΔminD2 in the presence and absence of plasmid pTA1392 from TEM analysed cells. n_Δ*pilB3*_ = 51, n_ΔminD2_ = 54, n_Δ*pilB3*_ = 51 n_Δ*pilB3* + pTA1392_ = 40 and n_Δ*minD2* + pTA1392_ = 59 (d) Semi-solid agar-based motility assay to visualise the swimming ability of H26 and Δ*minD2*. Left panel: representative inserts; right panel: average diameter of the motility rings. Graph represents values from 3 technical replicates from 3 different biological replicates. **** p<0.0001. Red arrow indicates the archaellum.

Given the observed effects of MinD2 deletion on archaellation levels, we investigated whether this impacted the motility of the Δ*minD2* strain, by growing the cells on soft agar plates. First, we found that Δ*minD2* strain grown in nutrient-rich YPC media showed reduced motility compared to H26 (data not shown). However, due to the composition of the medium, cell growth was rapid, making it difficult to differentiate growth from swimming halos in soft agar plates. Therefore, we switched to a different medium (CA), where strains grow slower. In order to support growth on CA medium, which lacks uracyl, strains were transformed with a plasmid (pTA1392) containing the *pyrE2* locus (which enables uracyl biosynthesis). Using this strategy, we observed that Δ*minD2* + pTA1392 cells exhibited a swimming defect, with their motility being only ∼25% of what was observed for H26 + pTA1392 cells (Figure 2d). We were able to complement the Δ*minD2* motility defect by expression of MinD2 on plasmid pTA1392 (pSVA6011, Figure 3). Additionally, the observed reduction in motility in Δ*minD2* was consistent with the lower number of mutant cells assembling archaella.

**Figure 3:**
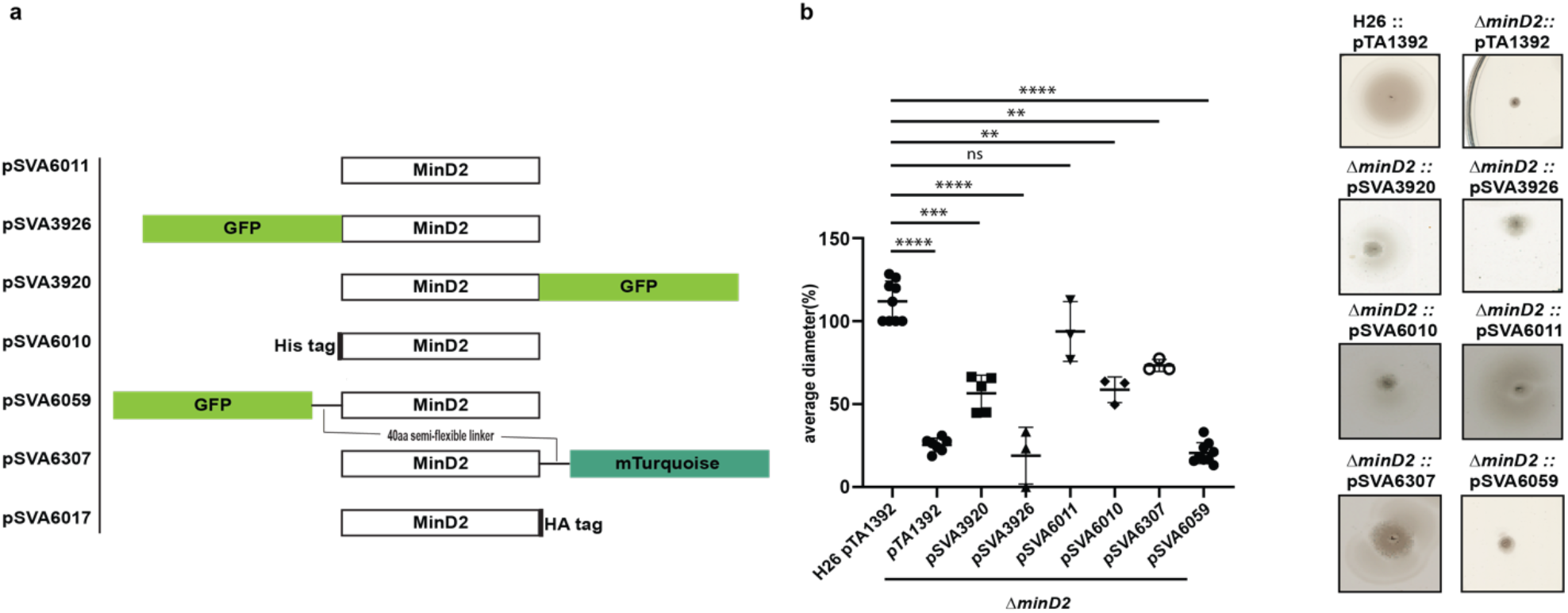
MinD2 plasmids and complementation assay. a) Schematic overview of MinD2 plasmids with different tags. b) Average diameter of motility rings measured relative to the wild type, from different *minD2* strains harbouring different tagged MinD2 plasmids. Values are from 3 independent experiments including 3 biological replicates each. (P values: **** <0.0001, *** 0.0002, ** 0.002, * 0.0332)

The hallmark of ParA/MinD superfamily proteins is the Walker A (WA) and Walker B (WB) motif (Leipe et al., 2002; Hu and Lutkenhaus, 2003). Mutation of both the motif in MinD4 had previously displayed motility phenotype and disrupted MinD4’s localisation at poles (Nußbaum et al., 2020). We wanted to investigate the role of the WA and WB motifs of MinD2 by generating plasmids with WA (K16A) mutant and WB (D117A). Expressing these in Δthe minD2 strain, we observed that mutation of WA did not have an effect on the swimming motility, and the strain had 100% activity as visualized in the wild type (H26). Whereas the WB mutant showed a defect in motility (∼30%) similar to Δthe *minD2* deletion mutant strain (Figure S4).

Thus, these results confirm that MinD2 not only modulates cells shape but also impacts cell motility, and that Walker B motif of MinD2 plays a role in the motility phenotype.

### Fluorescently tagged MinD2 shows diffused localisation

The observed impact of MinD2 on motility is reminiscent of the role of MinD4, which we previously showed to be a MinD homologue that oscillates along the cell axis in *H. volcanii* and stimulates the formation of chemosensory arrays and archaella at the cell poles, thereby regulating motility (Nußbaum et al., 2020). Therefore, we proceeded to examine the localisation pattern of MinD2, by generating fluorescent fusion proteins. For this, the MinD2 protein was tagged at the N- or C-terminus with various tags and linkers, which were then expressed in the Δ*minD2* deletion mutant. To test whether the fluorescent tag had an impact on the function of MinD2, we performed complementation experiments and assessed their motility using motility assays (Figure 3b). Notably, when a tag-less MinD2 variant was used (pSVA6011), the swimming phenotype could be restored to ∼91% of the activity observed in WT cells. Expression of an N-terminal His-tag MinD2 variant (pSVA6010) restored ∼62% of the motility activity, whereas the strain expressing an N-terminally tagged GFP variant (pSVA3926) had a motility defect similar to the Δ*minD2* strain (Figure 3b) and showed diffuse localisation in the cells (Figure S3). Expression of a variant in which MinD2 was fused to an N-terminal GFP tag with a semi-flexible linker (Ithurbide et al., 2024) (pSVA6059) could also not restore motility. Tagging MinD2 on the C-terminus was, in general, more effective at restoring motility. Using only an HA tag at the C-terminus complemented the swimming back to 100%

(Figure S4). And, the strain expressing a C-terminally GFP tagged MinD2 variant (pSVA3920) retained ∼60% of the motility displayed by WT cells, and the introduction of a semi-flexible linker between MinD2 and a C-terminal mTurquoise tag (pSVA6307) led to a restored swimming efficiency up to ∼80% of that observed in H26 cells (Figure 3, table 4).

We then proceeded to characterize the localization of the most functional C-term fluorescently tagged MinD2 variants. For this, samples were collected at similar OD^600^s as that of the cell shape experiments. Visualisation of both the C-terminally GFP tagged and C-terminally mTurquoise tagged MinD2 variants showed a diffused localisation throughout the cells (Figure S3b) and from early log phase through mid-log phase. Based on our observations, MinD2-linker-mturquoise didn’t show a change in localization during the transition from rod to plate phases and does not show a different localization in rods versus in plates. Collectively, these data show that MinD2 does not localize to the cell poles, suggesting that its impact on the polar organization of the motility machinery likely requires the interaction of MinD2 with other proteins. Furthermore, the inability of N-terminally tagged MinD2 to complement the swimming phenotype suggests that the N-terminus of MinD2 is important for its activity and interaction with protein partners in vivo.

### Possible interaction partners of MinD2

#### HVO_0596

HVO_0596 is a protein of unknown function encoded downstream of the *minD2* gene, likely within the same operon (Babski et al., 2016) (Figure 4a). Synteny report on MinD2 and HVO_0596 shows the two gene to be conserved in Haloarchaea (Table 5). Furthermore, an alphafold <u>3</u> (Abramson et al., 2024) prediction indicated a possible interaction between MinD2 and HVO_0596, based on 6 hydrogen bonds between 5 residues of HVO_0596 at the C-terminus and the C-terminus of MinD2 with an iPTM score = 0.83 and pTM = 0.63 (Figure 4b). To explore a potential role of HVO_0596 in MinD2 functionality, we generated a deletion mutant for HVO_0596 and a double deletion mutant *Δhvo_0596ΔminD2*. The *Δhvo_0596* deletion mutant showed no discernible impairment of cell growth (Figure 4c) and to the contrary of *minD2* deletion, *the deletion of HVO_0596 has no effect on* cell shape development and transition during growth (Figure 4d).

**Figure 4:**
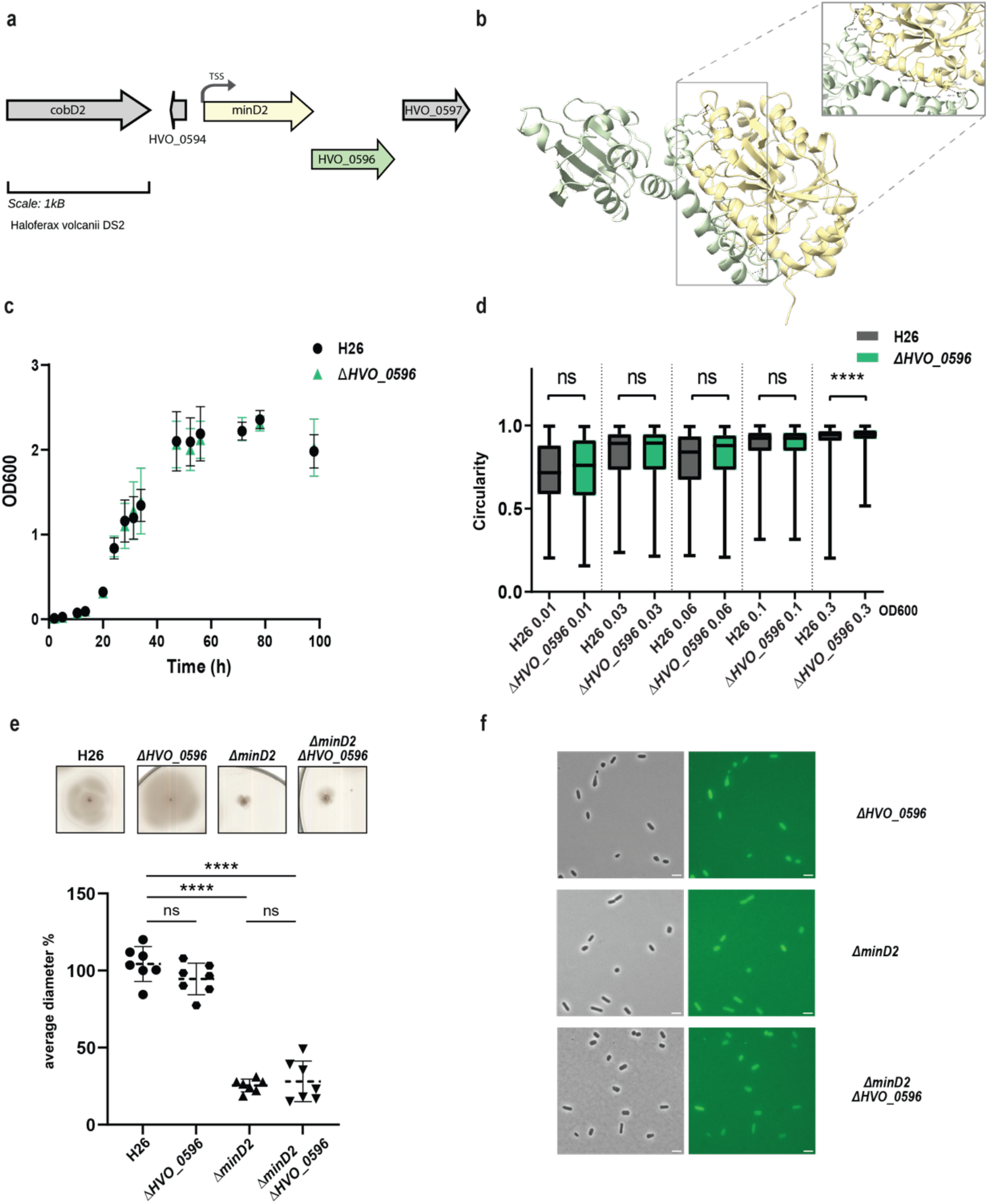
HVO_0596 might interacts with MinD2 but its deletion mutant has no phenotypes. (a) Schematic representation of the gene neighbourhood showing MinD2 (HVO_0595) (yellow) and HVO_0596 (green) showing the TSS (black arrow) present 28bp upstream of start. (b) Alphafold 3 prediction revealing interaction between MinD2 and HVO_0596. Black dashed lines indicates the intermolecular H-bond at the C-terminus of both proteins and red dashed line indicates the interaction found with relaxed angle criteria in ChimeraX. (c) Growth curve of H26 (black) and Δ*HVO_0596* (green) (d) Distribution of cell circularity (%) at different ODs; n >1100 (e) Quantification (bottom panel) of motility diameter for by the different mutants and WT (inserts: Top Panel). Calculations were made using 3 independent experiments including >2 biological replicates each. Black line indicates mean, lower, and upper lines the standard deviation. (f) Representative inserts of mNeonGreen-HVO_0596 localisation in H26(WT) and deletion mutants Δ*hvo_0596*, Δ*minD2* and Δ*HVO_0596minD2*. Scale bar: 4µm

Additionally, no soft-agar motility defect could be observed for the single mutant *Δhvo_0596 and* the double deletion mutant *Δhvo_0596ΔminD2* exhibited the same motility defect as the *ΔminD2* deletion mutant (Figure 4e). These results suggest that HVO_0596 does not directly affect motility.

To elucidate the localisation pattern of HVO_0596, we generated a fluorescently tagged version of HVO_0596 at its N-terminus with mNeongreen (pSVA6051). Localisation experiments showed diffuse fluorescence in both deletion mutants *Δhvo_0596, ΔminD2* and *Δhvo_0596ΔminD2* (Figure 4f).

#### CetZ5 and CetZ6

*H. volcanii* has 6 paralogues of CetZs of which CetZ1 and CetZ2 have been studied for their role as cytoskeletal proteins and role in motility (Brown et al., 2024). An accompanying study by (Brown et al, 2024, unpublished) suggests a possible interaction between MinD2 and CetZ1. Specifically, MinD2 was found to influence the cellular positioning of CetZ1, impacting its polar localisation. While the function of the other four CetZs is uncharacterised, CetZ5 was hypothesized to be a cytoskeletal protein by (Schiller et al., 2023). It is therefore possible that MinD2 interacts with these CetZ proteins in the cell. Therefore, we wanted to address possible interactions with CetZ5 and CetZ6.

CetZ5 and CetZ6, initially characterised as FtsZ7 and FtsZ8, belong to the tubulin/FtsZ family, and previous studies have shown that deleting these proteins has no effect on cell division (Duggin et al., 2015). To characterise the role of these proteins with respect to MinD2, we generated single deletion mutants (*ΔcetZ5* and *ΔcetZ6*) and double deletion mutants with *minD2* (*ΔcetZ5ΔminD2* and *ΔcetZ6ΔminD2*). We found no motility defects for either *ΔcetZ5* or *ΔcetZ6* (Figure 5a, 5c) as observed previously (Duggin et al., 2015). The double deletion mutants (*ΔcetZ5ΔminD2* and *ΔcetZ6ΔminD2*) showed reduced motility similar to that of the single *ΔminD2* mutant, indicating that the impact in swimming ability is due to deletion of MinD2 rather than these CetZ proteins.

**Figure 5:**
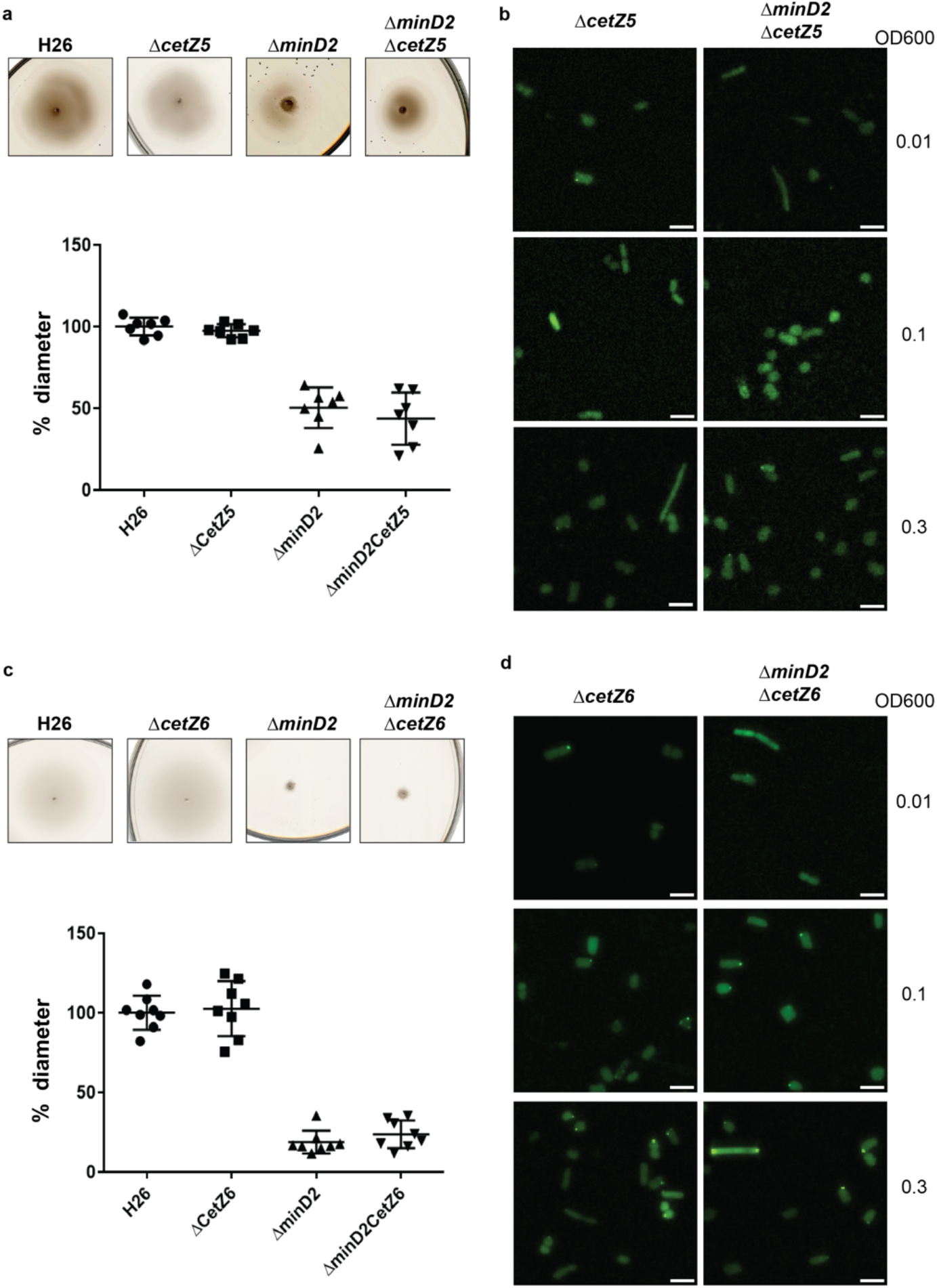
CetZ5 and CetZ6 do not have a motility phenotype and display diffused fluorescence. (a) Semi-solid agar-based motility assay for H26, Δ*CetZ5* and double mutant, Δ*minD2CetZ5*. (b) Localisation of GFP-CetZ5 in Δ*CetZ5* and Δ*minD2CetZ5*. (c) Semi-solid agar-based motility assay for H26, Δ*CetZ6* and double mutant, Δ*minD2CetZ6*. (b) Localisation of GFP-CetZ6 in Δ*CetZ6* and Δ*minD2CetZ6*. Scale bar: 4µm. Calculations were made using 3 independent experiments including 3 biological replicates each.

To gain insight into the cellular positioning of the CetZ proteins, we created N-terminally GFP tagged versions of CetZ5 (pSVA6040) and CetZ6 (pSVA6042). In both cases, the localisation of the proteins was diffused across the cells (Figure 5 c, 5d). To check if cell shape has an effect on CetZ localisation, we further visualized the distribution of the tagged proteins at different ODs. However, both GFP-CetZ5 and GFP-CetZ6 displayed diffused fluorescence throughout all the growth phases analyzed (Figure 5 c, 5d).

Collectively, our experiments indicate no direct interaction of HVO_0596, CetZ5, and CetZ6 with MinD2.

### MinD2 regulates the positioning of the motility and chemotactic machineries

Given that the *Δ*minD2 mutant exhibited a motility defect and displayed reduced assembly of archaella, we hypothesised that MinD2 may have a function similar to that of MinD4 with respect to the cellular positioning of the motility and chemotactic machineries. Therefore, we investigated the localization of the motility machinery in *Δ*minD2. For this, we used strains expressing ArlD-GFP, a fluorescently tagged version of a well-established marker for the archaellum motor complex, used as an indicator to identify cells with archaella (Li et al., 2019). Upon expression of ArlD-GFP in the Δ*arlD* strain at OD^600^ 0.01, we observed that most cells (77%) had fluorescent foci at the cell poles, with only a few cells displaying diffused fluorescence (23%) (Figure 6a, b), similar to results previously observed in *H. volcanii* (Li et al., 2019). However, when ArlD-GFP was expressed in *ΔminD2*Δ*arlD* cells, fluorescent foci at the poles were only detected in 25% of the cells (Figure 6a, b). This result is in agreement with the low abundance of archaella observed in TEM, where these structures were present in only ∼32% of *ΔminD2ΔpilB3* (+ pTA1392) cells (Figure 2).

**Figure 6:**
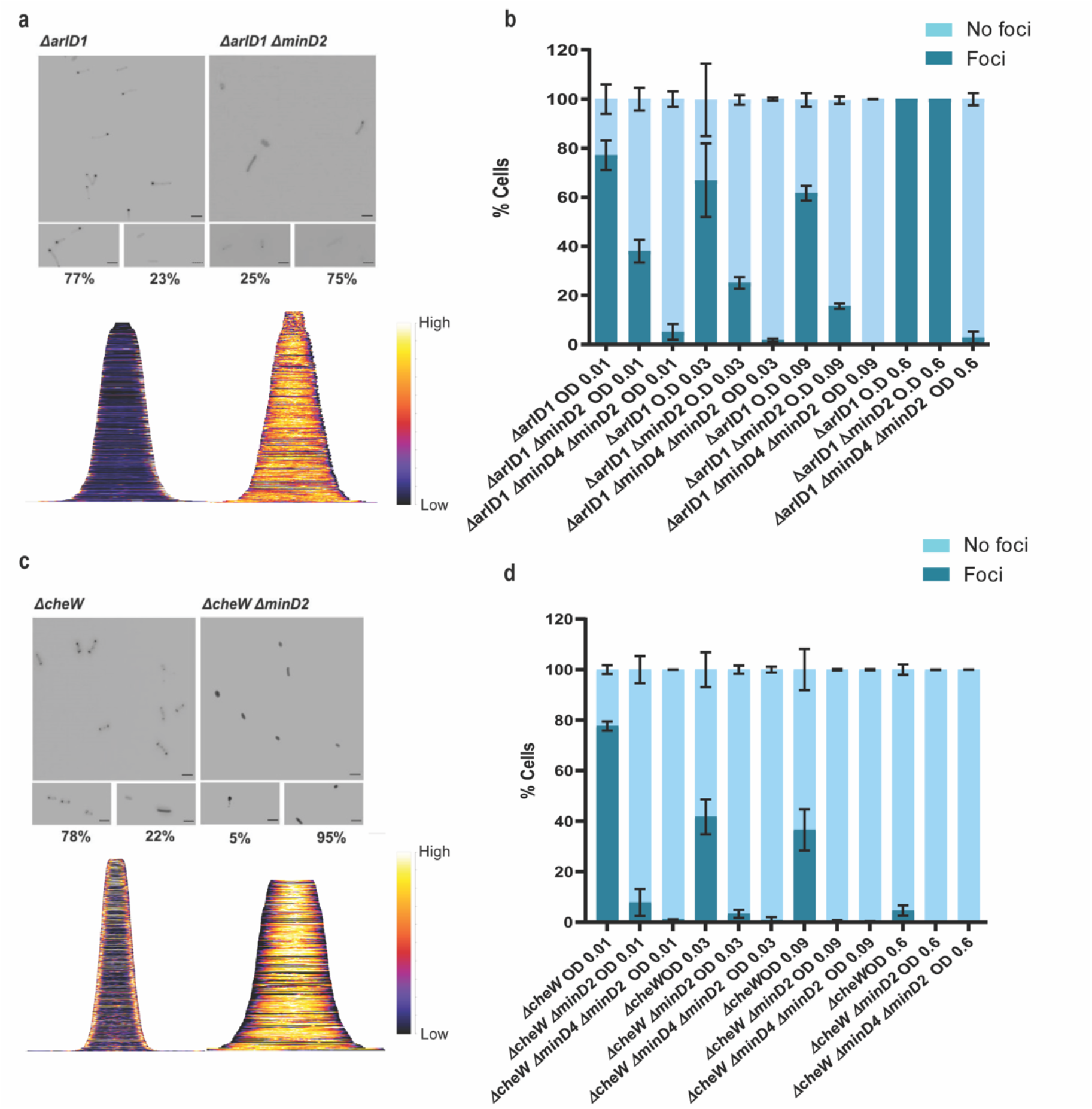
MinD2 affects the archaellum and chemotaxis machinery. (a) Fluorescent image of ArlD-GFP in Δ*arlD* and Δ*arlDminD2* strain. (b) Graphical analysis of the microscopic images to represent the % of cells with or without ArlD-GFP polar foci. (c) Fluorescent image of GFP-CheW in Δ*cheW* and Δ*cheWminD2* strain. (d) Graphical analysis of the microscopic images to represent the % of cells with or without GFP-CheW polar foci. Scale bar: 4µm. (a,c) Lower panel – Demographic analysis of the foci distribution showing spatial distribution of the proteins (yellow) arranged in an ascending order of cell length. Cells are arranged in ascending order (b,d). Calculations were made using 3 independent experiments including 3 biological replicates each.

As described above, in previous studies we established a correlation between MinD4 and the positioning of the archaellum machinery. Therefore, we decided to explore the links between MinD2 and MinD4 in the regulation of this process, by comparing the localization of the motility machinery in cells lacking either one of the MinD homologues (Δ*minD2*Δ*arlD* or ΔminD4Δ*arlD*) or both homologues (Δ*minD2*Δ*minD4*Δ*arlD*). In the Δ*minD4*Δ*arlD* mutant, we observed that the number of cells with ArlD polar foci was ∼20% at an OD^600^ of 0.01 (Nußbaum et al., 2020). In the *ΔminD2ΔminD4* mutant, the formation of polar foci decreased, being present in only ∼7% of cells at an OD^600^ of 0.01, and continuing to decrease with increasing OD^600^. Additionally, it was previously observed that all cells in the stationary phase form foci corresponding to the archaellum motor complex. We observed comparable results in the Δ*arlD* and *ΔminD2*Δ*arlD* mutants, with all cells having foci upon reaching the stationary phase at OD^600^ 0.6 (Figure 6b). However, in the *ΔminD2ΔminD4ΔarlD* mutants, we observed a consistent reduction in foci formation even at high OD^600^, which was not observed in the individual mutants (Figure 6b). which suggests a synergistic effect or partially redundant role of MinD2 and MinD4 in the archaellum polar assembly.

In *H. volcanii*, chemotaxis involves the assembly of chemosensory arrays, which are preferentially localized at the cell poles during the early log phase and become diffused as the cells enter stationary phase (Li et al., 2019). Therefore, we investigated whether MinD2 also influences the localization of chemosensory arrays, using the chemotaxis protein CheW as a marker for these clusters (Li et al., 2019). For this, we expressed GFP-CheW in cells lacking CheW (Δ*cheW*) and cells lacking both CheW and MinD2 (*ΔminD2ΔcheW*). In Δ*cheW* cells, expression of GFP-CheW led to polar foci in 78% of cells and diffused localisation in 22% of cells (Li et al., 2019 and Figure 6c). By contrast, in *ΔminD2ΔcheW* cells, expression of GFP-CheW led to only 5% of the cells displaying polar chemosensory foci, while the remaining 95% of *ΔminD2ΔcheW* cells showed diffused GFP-CheW localisation, suggesting the absence of chemosensory arrays localization (Figure 6c). Further analysis showed that the number of cells with CheW foci formation decreased substantially in Δ*cheW* as the OD increased, while the localisation of CheW reduces 5% - 0% in *ΔminD2ΔcheW* cells (Figure 6d).

Since previous studies have shown that MinD4 regulates the positioning of chemosensory arrays, we further investigated the links between MinD2 and MinD4 in modulating CheW localisation. For this, we created the triple deletion mutant Δ*minD2*Δ*minD4ΔcheW*, which has then been used to study the localization of GFP-CheW. Our results showed a further reduction in CheW foci formation in *ΔminD2ΔminD4ΔcheW* cells compared to *ΔminD2ΔcheW* cells, with only 2% of cells showing polar GFP-CheW foci at OD^600^ of 0.01. In the transition of OD^600^ from 0.01 to 0.03, we observed GFP-CheW foci formation to be completely absent, with no cells displaying CheW foci (Figure 6d).

Collectively, these data show that MinD2 impacts the localisation of both the motility and chemosensory machineries, in addition to the effect that MinD4 has on the localisation of these cellular components.

## Discussion

*H. volcanii* encodes four homologs of the MinD protein, which unlike their counterparts in bacterial cells, are not involved in cell division and do not regulate the localization of the divisome (Nußbaum et al., 2020). Previous studies have started to elucidate the functions of MinD proteins in *H. volcanii*, particularly MinD4 (HVO_0322), which governs the precise positioning of both the archaellum and chemotaxis machineries, which are indispensable for enabling directional and purposeful motility (Nußbaum et al., 2020). in rod cells. Here, we further extend the characterization of the functions of MinD proteins in *H. volcanii* cells, focusing on MinD2. Our analyses underscored MinD2 as a key player controlling cell shape morphology (Figure 1) and motility (Figure 2), by enabling cells to retain a rod shape, particularly in the early growth phase. Furthermore, we demonstrate that MinD2 shows diffuse localization across the cell (Figure 3) and likely interacts with a variety of partner proteins to mediate its effects (Figures 4, 5). By analysing the localization of a variety of proteins involved in the formation of the archaellum and chemotaxis complexes, we also reveal that MinD2 modulates the placement of the motility and chemotaxis machineries (Figure 6). Finally, we investigate the links between MinD2 and MinD4, showing that the two MinD homologues have synergistic roles in linking cell shape and motility in *H. volcanii* (Figure 6).

Studies on *H. volcanii* provide insights into the regulation of cell shape in response to environmental cues. The cells undergo remarkable transformations in cell shape during different growth phases and conditions. These changes, from rod-shaped to flat, polygonal pleomorphic disks (plate shaped), have been a subject of interest due to their potential roles in adaptation and survival strategies (Halim et al., 2017; Li et al., 2019; de Silva et al., 2021). However, our understanding of the molecular mechanisms controlling archaeal cell shape determination is still developing. Previous studies identified proteins such as CetZ1, LonB, ArtA, PssA, and PssD as important regulators of this process (Duggin et al., 2015; Halim et al., 2017; Ferrari et al., 2020; Brown and Duggin, 2023). More recently, cell-shape mutants lacking the ability to form plates have been studied, including DdfA (disk determining factor), which is likely involved in the signalling pathways that determine cell shape. Additionally, studies on RdfA (rod-determining factor) and Sph3 (SMC-like protein) indicate that these proteins are also involved in the signal cascade that potentially regulates cell shape (Schiller et al., 2023). However, how these and other cell-shape determinants interact with each other, and how their activity is regulated by environmental conditions, remains unclear.

Our findings identify MinD2 as another protein that regulates cell shape in *H. volcanii* cells (Figures 1 and S2). One of the most distinctive features of the *ΔminD2* mutant is its preponderance to form plate-shaped cells, including in the early log phase, highlighting the importance of MinD2 in maintaining the rod-shaped morphology characteristic of *H. volcanii* cells. Furthermore, while the presence of a plasmid delayed the rod-to-plate cell shape transition, particularly in early log phase, the majority of Δ*minD2* + pTA1392 cells still displayed plate shape as the optical density of the cultures increased (Figure 1b). Therefore, although the presence of plasmid can partially prevent the loss of rod shape in the Δ*minD2* mutant early on, the absence of MinD seems to dominate the phenotype, resulting in the majority of cells being plates.

While our findings demonstrate that MinD2 regulates cell shape, the mechanism by which MinD2 operates remains unclear. Notably, we find that MinD2 shows a diffuse localization pattern across the cell, rather than localizing to specific foci, such as the cell pole (Figure 3). Furthermore, our experiments with various variants of MinD2 suggest that its N-terminus is important for activity, potentially through interactions with protein partners (Figure 3). Indeed, we identified several potential interacting partners of MinD2, including HVO_0596, CetZ5, and CetZ6. However, mutants lacking HVO_0596, a protein transcribed from the same operon as MinD2, or either of the CetZs, showed no discernible phenotypes with regards to morphology (Figure 4 and 5). Furthermore, our experiments with single vs double mutants show that the observed impacts of motility in these mutants is primarily attributable to MinD2 (Figure 4 and 5).

While future studies are needed to further elucidate how MinD2 regulates morphology and motility in *H. volcanii*, our findings provide some insights into the links between these two cellular processes. For example, we found that discoid *ΔminD2* cells have a significant decrease in the number of archaella, which results in decreased motility (Figure 2). In addition, MinD2 deletion also impacted the positioning of the chemotaxis machinery(Figure 6). One possibility is that these changes in the localization of archaella and chemosensory arrays result from the observed changes in morphology in *ΔminD2* cells, particularly their inability to retain rod shape. This possibility is supported by studies in other microbial species linking cell shape and the special organization of motility machinery. For example, in *E. coli*, the distribution of chemosensory arrays has been shown to have a preference to the curved membrane (Strahl et al., 2015; Draper and Liphardt, 2017). Another possibility is that MinD2 participates in the spatial organization of the archaeal cell pole. For example, some proteins, like bacterial ParA/MinD homologs (such as FlhG and ParC) rely on polar landmark proteins for localisation (Lutkenhaus, 2012). In *H. volcanii*, the MinD4 homolog oscillates along the cell axis and is hypothesised to function similarly to polar landmark proteins, thereby influencing the proper positioning of both archaella and chemosensory arrays (Nußbaum et al., 2020). Notably, here we show that *ΔminD2ΔminD4* mutants have stronger defects in the positioning of the motility and chemosensory machinery (Figure 6) than *ΔminD4* single mutants (Nußbaum et al., 2020 – Figure 6), suggesting that MinD2 and MinD4 have non-redundant roles in these processes.

Based on our findings and previous studies, we propose a model for how MinD2 controls cell shape and motility in *H. volcanii* (Figure 7). In this model, while MinD4 is important for promoting the adequate positioning of polar proteins, some of these proteins can still find the ‘probable’ pole even in the absence of MinD4. This is in agreement with previous studies, which showed that while a reduced frequency of *ΔminD4* mutant cells express archaella (20%) and chemosensory array (40%), some cells can still assemble these structures (Nußbaum et al., 2020). In the case of *ΔminD2* mutants, we postulate that the loss of rod shape resulting from MinD2 absence influences the positioning of these machineries in most cells, although some can still assemble archaella and chemosensory arrays guided by polar cap proteins. In the absence of both MinD2 and MinD4, cells are always plate shaped and polar MinD4 patches are not established. This combination may render proteins targeted to the poles unable to detect ‘probable’ poles and/or impair their interaction with different polar cap proteins. Under this scenario, the lack of both MinD2 and MinD4 leads to a synergistic effect, resulting in an almost complete absence of cells with archaellum (5%) or chemosensory (2%) machineries.

**Figure 7:**
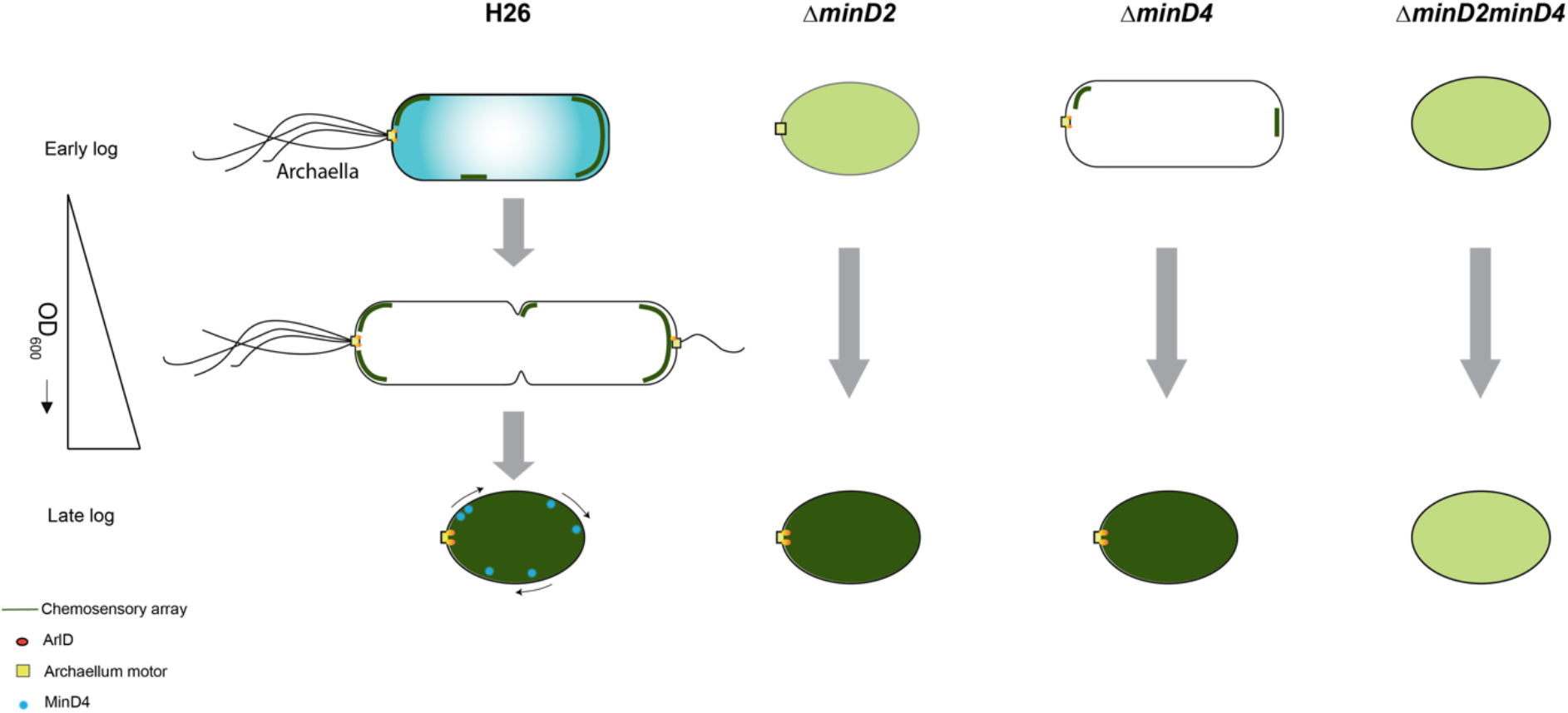
Proposed model for MinD2 function in *Haloferax volcanii*. *H. volcanii* cells show growth-dependent cell shape morphology transitioning from rods to plate-shaped cells. We show that MinD2 has a strong effect on shape, with Δ*minD2* mutant cells predominantly adopting a plate shape. We propose that the shape phenotype impacts the proper positioning of the archaellum (archaellum motor: yellow square and ArlD: orange ovals) and chemotactic machinery (dark green lines). According to this model, in Δ*minD2* cells, the localization of both machineries is diffused (light green) with a very few cells showing the ablility to localise the chemosensory arrays in early log phase. As the growth stage progresses, only chemosensory arrays remain diffused (dark green) and the archaellum motor is able to localise at a pole. Previous studies on MinD4 (Nußbaum et al., 2020), shows the Δ*minD4* mutant cells has an effect on the positioning on archaellum and chemosensory arrays. Together, we postulate that deleting both MinD homologues (Δ*minD2minD4*) has a synergistic effect, resulting in these cells being unable to localize both archaellum and chemosensory arrays to the pole at any stage of the growth phase.

Morphological integrity and the determination of cell shape are pivotal aspects of microbial physiology. While these mechanisms are fairly well understood in bacteria, less is known about their archaeal counterparts. This study further advances our understanding of cell morphology determination in archaea by highlighting the pivotal role of the archaeal MinD2 homologue in modulating *H. volcanii* morphology. Notably, while bacterial Min proteins predominantly influence cell division, our observations suggest that MinD2 in archaea may have evolved or diversified to have more influence on cell morphology. Additionally, our findings support a model in which MinD2 contributes to cells retaining a rod-shape morphology, which enables polar cap proteins to recognise the cell poles, thereby supporting the assembly and adequate positioning of the motility and chemotaxis machinery. Furthermore, our findings add support to previous studies on MinD4 suggesting that archaeal MinD homologues have non-redundant roles in influencing cell shape and the positioning of motility and chemotaxis machinery. These findings contribute to a deeper understanding of the intricate regulatory network governing cellular processes in archaea, and set the stage for future studies aimed at uncovering the detailed molecular mechanisms by which MinD proteins modulate haloarchaeal physiology.

## Materials and Methods

All chemicals have been purchased from Roth or Sigma unless stated otherwise.

### Strain and growth condition

*E. coli* strains were cultured in LB (Luria Broth)-medium or grown on LB agar plates, with the necessary antibiotics (100 µg/mL ampicillin, 30 µg/mL chloramphenicol, 25 µg/mL kanamycin) and grown at 37° C. Liquid cultures were constantly shaken at 150 rpm.

*H. volcanii* H26 cells were grown in YPC medium [0.5%(w/v) yeast extract (Difco), 0.1%(w/v) peptone (Oxoid), and 0.1%(w/v) casamino acids (Difco)] dissolved in 18% buffered Salt Water (SW) (144 g/l NaCl, 18 g/l MgCl2 * 6 H2O, 21 g/l MgSO4 * 7 H2O, 4.2 g/l KCl, 12 mM Tris/HCl, pH 7.5), supplemented with 3mM CaCl^2,^ adjusted to a pH of 7.2 with KOH for transformations. For experiments, CAB medium [CA medium (0.5% (w/v) casamino acids (Difco)] dissolved in 18% SW, supplemented with 3mM CaCl^2^, and 0.8 µg/mL of thiamine, and 0.1 µg/mL of biotin, adjusted to a pH of 7.2 with KOH) supplemented with trace elements solutions (Duggin et al., 2015)] was used. For each experiment, a single colony was inoculated into 5mL medium and diluted to a larger volume on the subsequent day. This dilution was crucial to ensure an appropriate cell density for subsequent experiments. By adjusting the OD^600^ to the desired value, it was possible to obtain a consistent starting point on the day of the experiment.

For strains with an auxotrophic mutation grown in CA/CAB medium, the medium was supplemented with 50 µg/mL uracil for *ΔpyrE2*. Alternatively, the strains were transformed with a plasmid carrying the respective gene for viable growth. For growth curve, cells were grown in 15mL culture volume and measured with cell growth quantifier (CGQ) (Aquila biolabs GmbH) at 45°C and shaking at 120 rpm.

### Genetic modification of *Haloferax volcanii*

Transformation in H26 based on uracil selection via the Polyethylene glycol 600 (PEG600) along with gene deletion and expression studies were conducted as described previously (Allers et al., 2004). For transformation into *H. volcanii*, non-methylated plasmids were extracted from *E. coli dam*^*-*^*/dcm*^*-*^ (C2925I, NEB). Mutant strains generated and used are described in Table S1. Plasmids created for knockout mutants are described in Table S4. Primers to create knockout plasmids were based on pTA131, are described in Table S3.

### Growth Curve

Glycerol stocks were streaked on solid agar medium substituted with uracil and incubated at 45°C for 5 days. A single colony from plates was used to inoculate 5 mL of media on day 1. Strains without plasmid were grown in CAB with 50µg/mL uracil and strains with plasmid pTA1392 (containing p*fdx -pyrE2*) were grown in CAB medium without supplements. To generate a growth curve, the obtained culture was inoculated on day 2 at a starting OD^600^ of 0.05. The cell density was measured with cell growth quantifier (CGQ) (Aquila biolabs GmbH) at 45°C and shaking at 120 rpm and measurement were taken every 300s.

### Microscopy

The cell shape was analysed by imaging the cells with an inverted phase contrast light microscope (Zeiss Axio Observer Z.1). The cells were grown in 5mL of the respective medium and diluted in 20mL media volume the next day in order to achieve an OD^600^ of 0.01 the day after. For each culture, 5 µL sample was collected from different growth phases and dropped at the centre of an agarose pad (0.3% (w/v) agarose dissolved in 18% SW). On drying, the pad was covered with a cover slip and imaged. The images were acquired at 100x magnification using the oil immersion phase contrast (PH3) channel. All sampled were analysed in triplicated. Fluorescence microscopy images were acquired on Zeiss Axio Observer Z.1 (ex: 450-490 nm em: 500-550 nm filter from Chroma®), equipped with a heated XL-5 2000 Incubator running VisiVIEW℗ software for MinD4 and CheW.

### Image analysis

The phase contrast images from the microscopy were analysed using Fiji (Schindelin et al., 2012) combined with MicrobeJ plugin (Ducret et al., 2016). For the analysis, cells that formed aggregates or were fragmented were discarded from the calculation. The circularity of the cells was automatically calculated. The diameter of each analysed cell was thereon calculated and grouped into 6 bins in the range interval of 0.1 to 1. The parameters used for circularity were as previously defined in Patro et al., 2023.

### Transmission electron microscopy

Cells were harvested at 2000g for 15 mins. The resulting pellet was resuspended to a theoretical O.D of 10. 5µL of cells of the cell suspension was applied to a glow discharged carbon coated copper grid (Plano GmbH, Wetzlar Germany) and incubated for 10 seconds. The excess liquid was blotted away. The cells were then stained with 2% uranyl acetate (w/v). Cells were imaged using Zeiss Leo 912 Omega (tungsten) operated at 80kV and images were taken using Dual speed 2K on Axis charge-coupled device (COD) camera (TRS, Sharp-Eye).

### Motility Assay

Semi-solid agar plates were prepared using 0.33% agar in CA medium supplemented with 1 mM tryptophan. Cultures grown at OD^600^ 0.3 were inoculated into the plates using stab techniques, and the plates were then incubated at 45°C for 4 days. To compare the motility of different strains, all strains were spotted on the same plate. For each strain, a minimum of 3 technical replicates and 3 biological replicates were conducted. After 4 days, the diameter of the motility ring was assessed

### Western Blot

To assess the stability and proper expression of fusion proteins, similar cell cultures used for fluorescence microscopy analysis were collected. The cultures were harvested by centrifugation at 5000 rpm for 20 minutes, and the resulting pellet was resuspended in 1x SDS buffer containing PBS. Samples were loaded in a 11% SDS-PAGE gel and subsequently transferred to a PVDF membrane.

To block nonspecific binding, the membrane was treated with 0.1% I-BlockTM (Applied Biosystems, California USA) for 1 hour at room temperature. The membrane was then incubated with a primary antibody, [GFP (Sigma-aldrich, California USA) diluted at 1:5000 or anti-HA at a dilution of 1:10,000] for 3-4 hours at 4°C. For detection, a secondary antibody, anti-rabbit conjugated to Horse Radish Peroxidase (from goat) (ThermoFischer Scientific, Massachusetts USA), diluted at 1:10,000, was applied to the membrane and incubated for 1 hour at 4°C. The resulting image was developed using the Invitrogen TM I-BrightTM scanner (ThermoFischer Scientific).

### Structural analyses

The Alphafold models for MinD2 and HVO_0596 were generated with Alphafold 3 Google colab (Abramson et al., 2024)and further analyzed by ChimeraX (Meng et al., 2023).

## Supplementary information

Avalaible in the Supplementary Material file.

## Acknowledgments

MP was supported by the German Research Foundation on grant number AL1206/4-3. SG was supported by the German Science Foundation (DFG) project number 505545313 (AL1206/14-1) to SVA. PN and SI were supported by the VW Foundation by a Momentum grant to SVA (AZ 94993). SS was supported by the German Research Foundation under project number 403222702-SFB 1381 to SVA. We would like to thank the EM facility at the University of Freiburg for access to the TEM. The TEM (Hitachi HT7800) was funded by the DFG project number 42689454 and is operated by the faculty of biology at the University of Freiburg as a partner unit within the Microscopy and Image Analysis Platform (MIAP) and Life Imaging Centre (LIC), Freiburg. Molecular graphics and analyses performed with UCSF ChimeraX, developed by the Resource for Biocomputing, Visualization, and Informatics at the University of California, San Francisco, with support from National Institutes of Health R01-GM129325 and the Office of Cyber Infrastructure and Computational Biology, National Institute of Allergy and Infectious Diseases.

## Author contributions

MP and SVA conceived the experiments. Most of the experiments were performed by MP with the following exceptions; SS and MRF performed TEM. SG performed the WA and WB motility experiments and analysis. PN and SI constructed plasmids pSVA6059 and pSVA6307, respectively. SI and SVA supervised MP. MP and SVA wrote the manuscript. All authors checked the manuscript.

