## Supplementary Material for "MinD2 modulates cell shape and motility in the archaeon *Haloferax volcanii*"

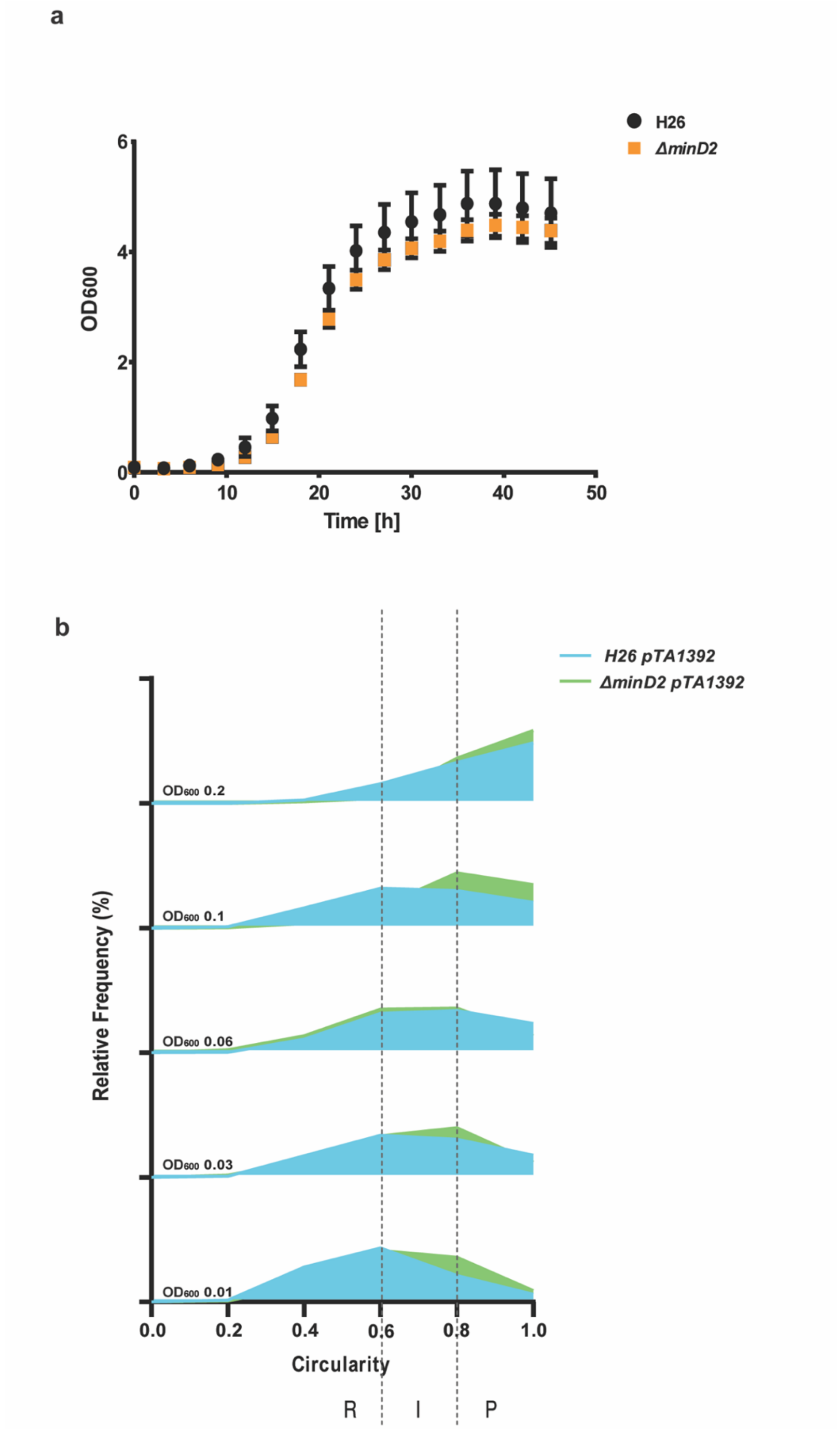


**Supplementary figure 1:** (**a**) Growth curve measurement of H26 and *ΔminD2* shows no growth defect in the mutants. (**b**) Relative Frequency measurement of cell circularity of H26 and *ΔminD2* in the presence of plasmid pTA1392. Vertical dashed line represents the different cell type R: Rods, I: Intermediates and P: Plates. Sum of the graph height per OD$_{600}$ equals 100% and Y-axis indicates the percentage of cell population per cell type

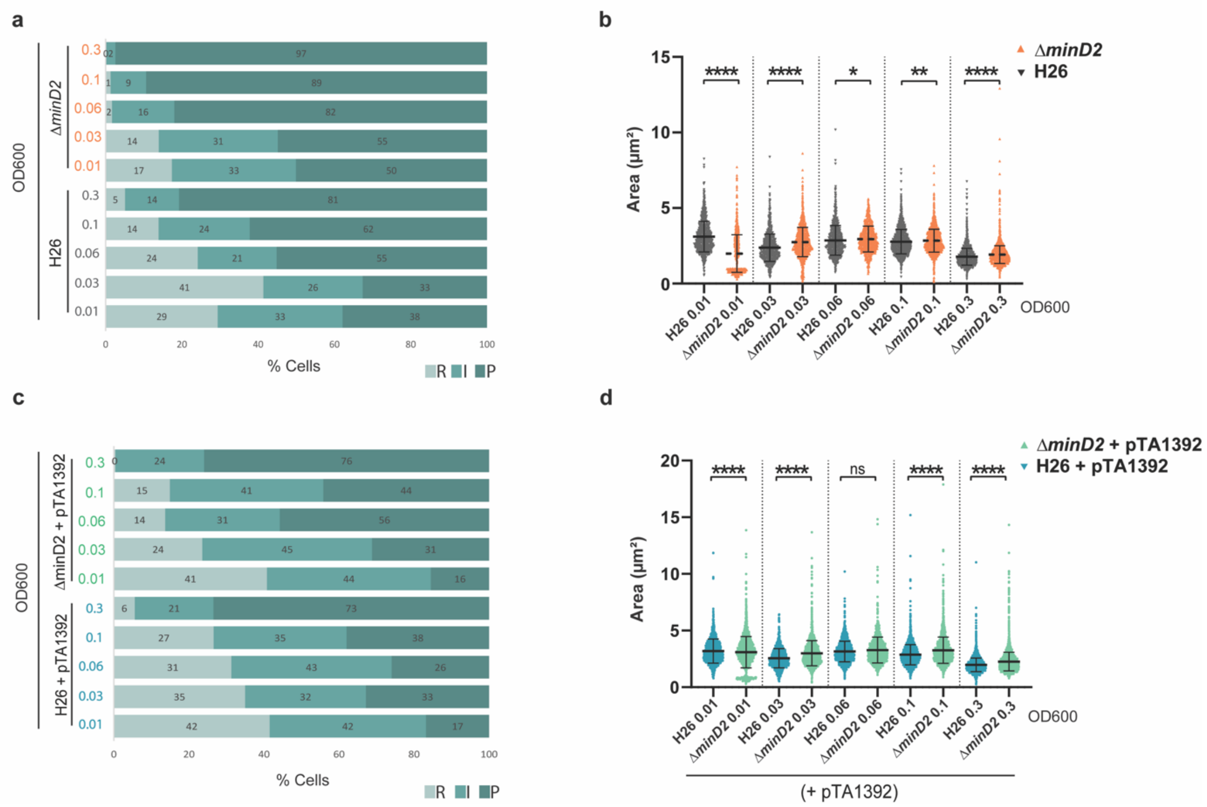


**Supplementary figure 2: Morphological analysis of strains and plasmid**. Bar graph showing the percentage cell present in each cell type (Rods (R), Intermediates (I) and Plates(P)) for (**a**) H26 (grey) and *ΔminD2* (orange); and (**c**) H26 +pTA1392 (blue) and *ΔminD2* + pTA1392 (green). (**b, d**) Scatter plot distribution of cell area (µm$^2$) at different $OD_{600}$. $n_{H26}$ >1363 n $_{\Delta minD2}$ > 2769. Calculations were made using 3 independent experiments including more than 3 biological replicates each.

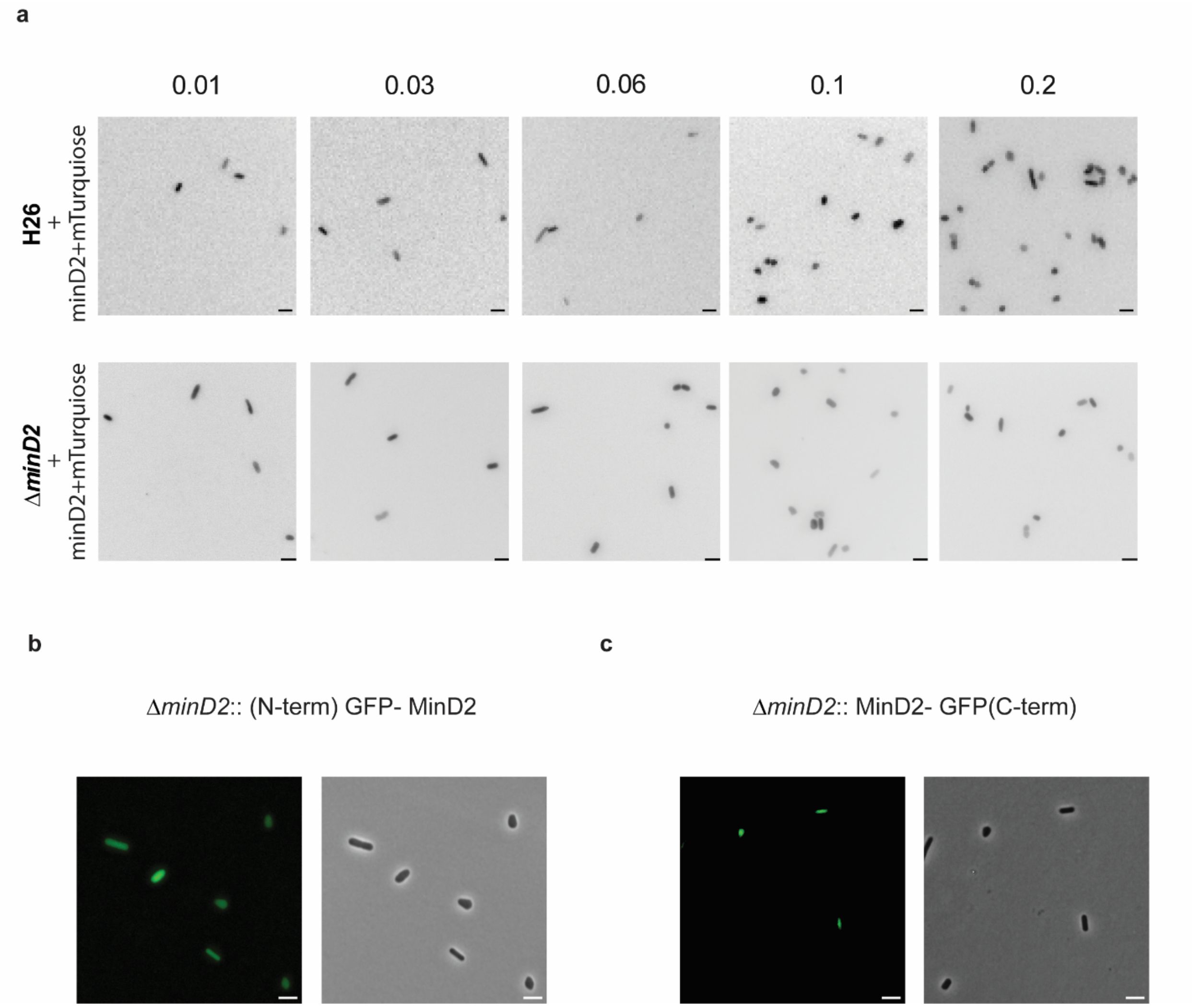


**Supplementary figure 3: Localisation of MinD2 shows diffused fluorescence** (**a**) MinD2 + semi flexible linker + mTurquoise showing diffused fluorescence in both wild type (top panel) and *ΔminD2*(bottom panel). Diffused GFP localisation for (**b**) N-term tagged GFP MinD2 and (**c**) C-term tagged MinD2 GFP; Insert are taken at OD 0.06. Scale bar = 4μm

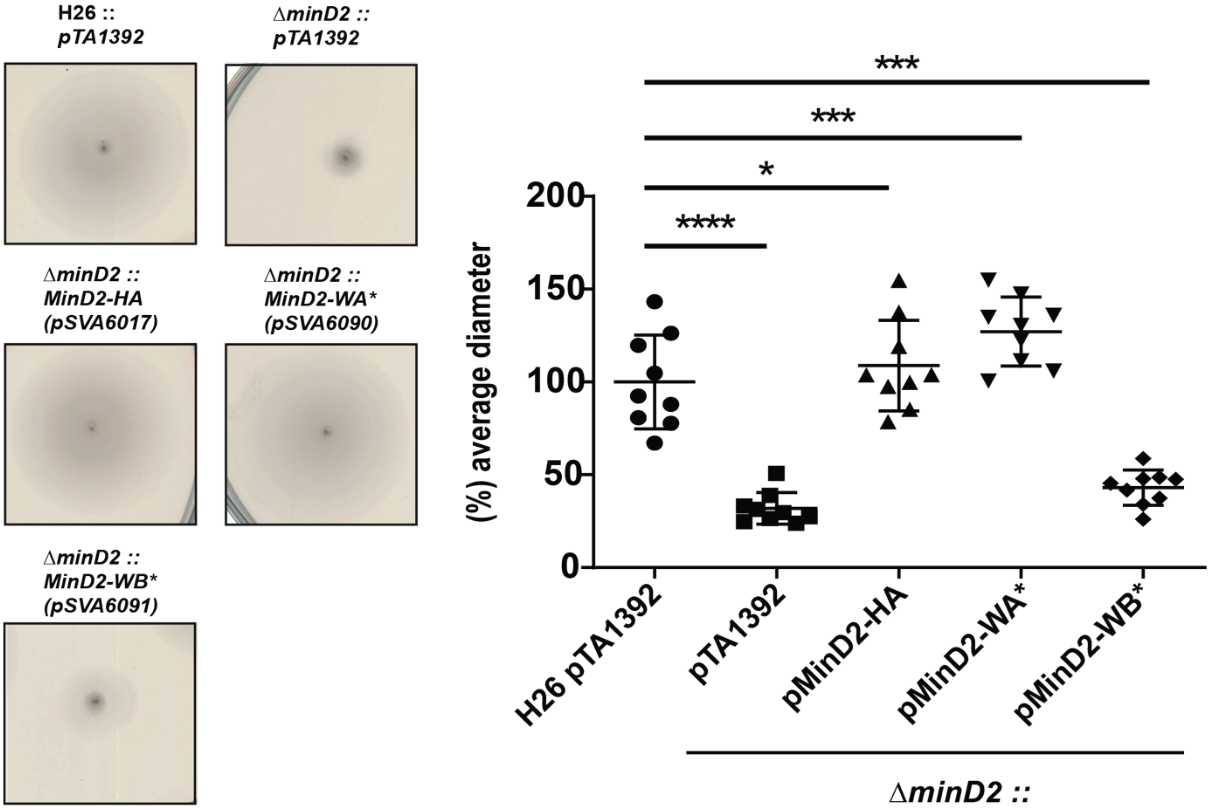


**Supplementary figure 4 Complementation of WA* and WB* in MinD2** (**a**) Motility assays to test swimming activity of WA and WB mutant in *ΔminD2* strain. pMinD2-HA was used as positive control as it was shown to complement full motility function in *ΔminD2* strain.

**Table 1: Strains used in this study**

| **Strain name** | **Genotype** | **Reference** |
|---|---|---|
| ***H.volcanii*** | | |
| H26 | *ΔpyrE2* | (Allers et al., 2004) |
| HTQ19 | *ΔpyrE2ΔflaD1* | (Li et al., 2019) |
| HTQ228 | *ΔpyrE2ΔminD2* | (Nußbaum et al., 2020) |
| HTQ241 | *ΔpyrE2ΔminD4ΔminD2* | (Nußbaum et al., 2020) |
| HTQ256 | *ΔpyrE2ΔflaD1ΔminD2* | This study |

| | | |
|---|---|---|
| HTQ255 | Δ*pyrE2*Δ*flaD1*Δ*minD2*Δ*minD4* | This study |
| HTQ247 | Δ*pyrE2* Δ*pilB3* | (Nußbaum et al., 2020) |
| HTQ248 | Δ*pyrE2*Δ*pilB3*Δ*minD2* | (Nußbaum et al., 2020) |
| HTQ451 | Δ*pyrE2*Δ*cheW*Δ*minD2* | This study |
| HTQ452 | Δ*pyrE2*Δ*HVO_0596* | This study |
| HTQ453 | Δ*pyrE2*Δ*cheW*Δ*minD2*Δ*minD4* | This study |
| HTQ454 | Δ*pyrE2*Δ*cetZ5* | This study |
| HTQ456 | Δ*pyrE2*Δ*cetZ6* | This study |
| HTQ457 | Δ*pyrE2*Δ*cetZ5*Δ*cetZ6* | This study |
| HTQ460 | Δ*pyrE2*Δ*minD2*Δ*HVO_0596* | This study |
| ***E.coli*** | | |
| 10-beta Competent Cells"TOP10" | Δ*(ara-leu) 7697 araD139 fhuA* Δ*lacX74galK16 galE15 e14-* ϕ*80*d*lacZ*Δ*M15 recA1 relA1 endA1 nupG rpsL* ($Str^R$) *rph spoT1* Δ*(mrr-hsdRMS-mcrBC)* | New England Biolabs |
| *$dam^-$/$dcm^-$* Competent cells | *ara-14 leuB6 fhuA31 lacY1 tsx78 glnV44 galK2 galT22 mcrA dcm-6 hisG4rfbD1 (zgb210::Tn10)* $Tet^S$ *endA1 rspL136* ($Str^R$) *dam13::Tn9* ($Cam^R$) *xylA-5 mtl-1 thi-1 crB1 hsdR2* | New England Biolabs |

**Table 2: plasmid used in this study**

| Plasmids | Description | Primer used | Enzyme used | Source/reference |
|---|---|---|---|---|
| pTA131 | Integrative plasmid with a *pyrE2* selection marker for knock-outs in *H. volcanii* ($Amp^r$) | - | - | (Allers et al., 2004) |
| pTA1392 | Overexpression plasmid for *H. volcanii* under the control of a tryptophan | | | (Gamble-Milner R, 2016) |

| | | | | |
|---|---|---|---|---|
| | inducible promotor. Contains *pyrE2* selection marker(Amp$^r$) | - | - | |
| pIDJL40 | Plasmid to express proteins with a C-terminal gfp phusion under the control of a tryptophan inducible promotor. Contains *pyrE2* selection marker (Amp$^r$) | - | - | (Duggin et al., 2015) |
| pHVID21 | Plasmid to express proteins with a C-terminal mTurquoise under the control of a tryptophan inducible promotor. Contains *pyrE2* selection marker (Amp$^r$) | - | - | Duggin lab |
| pSVA1841 | Integrative plasmid with a *pyrE2* selection marker to knock-out *hvo_0595* (*minD2*)(Amp$^r$) | - | - | (Nußbaum et al., 2020) |
| pTQ99 | Integrative plasmid with a *pyrE2* selection marker to knock-out *hvo_1200* (flaD1/arlD1)(Amp$^r$) | - | - | (Li et al., 2019) |
| pSVA5029 | Integrative plasmid with a *pyrE2* selection marker to knock-out *hvo_1225* (cheW)(Amp$^r$) | - | - | (Li et al., 2019) |
| pSVA5993 | Integrative plasmid with a *pyrE2* selection marker to knock-out *hvo_0596* (Amp$^r$) | 11068-11071 | - | This study |
| pSVA6037 | Integrative plasmid with a *pyrE2* selection marker to knock-out *hvo_2013 (CetZ5)* (Amp$^r$) | 10651-10654 | - | This study |
| pSVA6038 | Integrative plasmid with a *pyrE2* selection marker to knock-out *hvo_2068 (CetZ6)* (Amp$^r$) | 10655-10658 | - | This study |
| pSVA6039 | Integrative plasmid with a *pyrE2* selection marker for double knock-out *hvo_0595 and hvo_0596 (minD2 and HVO_0596)* (Amp$^r$) | 6919,6920, 10659,10660 | - | This study |
| pSVA3919 | Plasmid to express *flaD1* with a C- terminal gfp-tag(Amp$^r$) | - | - | (Li et al., 2019) |
| pSVA5031 | Expression plasmid for CheW-GFP | - | - | (Li et al., 2019) |
| pSVA3920 | Expression plasmid for MinD2 with a C-terminal GFP | 6575,6576 | NdeI, BamHI | This study |

| | | | | |
|---|---|---|---|---|
| pSVA3926 | Expression plasmid for N-terminal GFP with MinD2 | 8009, 8010 | NheI, BamHI | This study |
| pSVA6010 | Expression plasmid for N-terminal His tag-MinD2 | 10605, 10606 | NcoI, EcoRI | This study |
| pSVA6011 | Expression plasmid for tag-less MinD2 | 6575, 10606 | NdeI, EcoRI | This study |
| pSVA6059 | Expression plasmid for a N-terminal GFP-Linker-MinD2 | 10697, 10698 | NheI,NheI | This study |
| pSVA6017 | Expression plasmid for MinD2with C-terminal HA tag for IP experiment | 10605,10610 10614,10615 | NcoI, EcoRI | This study |
| pSVA6307 | Expression plasmid for MinD2 with a C-terminal mTurquoise | 6575, 10628 | NdeI, BamHI | This study |
| pSVA6040 | Expression plasmid for N-terminally tagged HA-GFP-CetZ5 | 10665, 10666 | NheI, BamHI | This study |
| pSVA6042 | Expression plasmid for N-terminally tagged HA_GFP-CetZ6 | 10669, 10670 | NheI, BamHI | This study |
| pSVA6051 | Expression plasmid for n-terminal taggedmNeonGreen-HVO_0596 | 10647, 10648 | NheI, BamHI | This study |

**Table 3: Primer used in this study**

| Primer number | Sequence | Description |
|---|---|---|

| | | |
|---|---|---|
| 6575 | GTTCTACATATGGTCGAGGCGTTC GCCGTCGCCAG | Forward primer for MinD2 with C-terminal GFP tag with NdeI restriction site |
| 6576 | GTAGGATCCCTCGGGGACGACGG CGCTTTTG | Reverse primer for MinD2 with C-terminal GFP tag with BamHI restriction site |
| 8009 | TCTAGCTAGCGAGGCGTTCGCCGT CGCCAG | Forward primer for MinD2 with N-terminal GFP tag with NheI restriction site |
| 8010 | CTAGGATCCGGGATTCTCATATTC GCTC | Reverse primer for MinD2 with N-terminal GFP tag with BamHI restriction site |
| 10605 | AGGACCATGGTCGAGGCGTTCGC | Forward primer for MinD2 with NcoI restriction site |
| 10606 | CGCGAATTCTCATATTCGCTCG | Reverse primer for MinD2 with EcoRI restriction site |
| 10610 | TAAGCGGGAATTCTCACGCGTAGT CCGGGACGTCGTACGGGTAGCTG CCTATTCGCTCGGGGA | Forward primer to insert MinD2 for C-terminal HA tag with EcoRI restriction site |
| 10628 | GAACGGATCCTATTCGCTCGGGGA CGACGG | Forward primer to amplify minD2 with BamHI restriction site |
| 10647 | AGGTGGCTAGCAGAATCCCGCGG GGCGAA | Forward primer to insert HVO_0596 with NheI restriction site |
| 10648 | ACCTGGATCCTCAGCGACTGTCCG GGCCGGC | Reverse primer to insert HVO_0596 with BamHI restriction site |

| | | |
|---|---|---|
| 10651 | GGCGAATTGGGTACCTTCCCGACGACCGACGACTG | Forward primer for US gene amplification for CetZ5 deletion-overlap PCR (orange represent overlap to plasmid pTA131) |
| 10652 | CCCGACCGCCTCGACGATAGCTCCCG | Reverse primer for US gene amplification for CetZ5 deletion- overlap PCR |
| 10653 | GTCGAGGCGGTCGGGCGTCTCTCTTTGACACGTC | Forward primer for DS gene amplification for CetZ5 deletion- overlap PCR (blue overlap US region) |
| 10654 | GGCGGCCGCTCTAGATCACTCGTCGCCCGCGCG | Reverse primer for DS gene amplification for CetZ5 deletion- overlap PCR (orange represent overlap to plasmid pTA131) |
| 10655 | GGCGAATTGGGTACCACTCAGCGGTAAATCCGATC | Forward primer for US gene amplification for CetZ6 deletion- overlap PCR (orange represent overlap to plasmid pTA131) |
| 10656 | ACGCACGGGGGTCGATAAACGTCGC | Reverse primer for US gene amplification for CetZ6 deletion- overlap PCR |
| 10657 | TCGACCCCCGTGCGTGTTGTCTGCTCTGCGACGTACC | Forward primer for DS gene amplification for CetZ5 deletion- overlap PCR (blue overlap US region) |
| 10658 | GGCGGCCGCTCTAGATCGTGCTCGCGCTCGGCGGC | Reverse primer for DS gene amplification for CetZ6 deletion- overlap PCR (orange represent overlap to plasmid pTA131) |

| | | |
|---|---|---|
| 10659 | ATTACCATATGCACGCTTCTCGAC GGTTGAG | Forward primer for double deletion of minD2 and HVO_0596 with NdeI restriction site |
| 10660 | ATTACCATATGCACGCTTCTCGAC GGTTGAG | Reverse primer for double deletion of minD2 and HVO_0596 with XbaI restriction site |
| 10665 | ATTACCATATGCACGCTTCTCGAC GGTTGAG | Forward primer to amplify CetZ5 with nheI restriction site |
| 10666 | TatgGGATCCTCAGAACAGCGAGTC GAGGC | Reverse primer for CetZ5 amlification with BamHI site |
| 10669 | TACCGCTAGCAACGTGTTCTGCTT TGG | Forward primer to amplify CetZ6 with NheI restriction site |
| 10670 | attgGGATCCTCACGCGTCGCCCGA GTCAC | Reverse primer for CetZ6 amlification with BamHI site |
| 10697 | CGCGCTAGCCTTGAGGGTAGCGG ACAAGG | Forward primer to incorporate semi-flexible Linker (from Alex Bisson) |
| 10698 | CGCGCTAGCGCCTTGACCTGGGC CAGATC | Reverse primer to incorporate semi flexible Linker |
| 11068 | GGCGAATTGGGTACCACTGCGAAA GCGAACGATTG | Forward primer to amplify US gene for deletion of Hvo_0596. Overlap PCR |

| | | |
|---|---|---|
| 11069 | CAACCGTCGAGAAGCGTGTCGCC CCGCGGGATTC | Reverse primer to amplify US gene for deletion of Hvo_0596. Overlap PCR |
| 11070 | GAATCCCGCGGGGCGACACGCTT CTCGACGGTTGAG | Forward primer to amplify DS gene for deletion of Hvo_0596. Overlap PCR |
| 11071 | GGCGGCCGCTCTAGACCAGTCCG CGAAGTCGG | Reverse primer to amplify DS gene for deletion of Hvo_0596. Overlap PCR |

**Table 4: Swimming motility of various mutants used in this study.** Average diameter of motility rings measured relative to the wild type, from different strains.

| Strain | Average motility diameter (%) |
|---|---|
| H26 pTA1392 | 100 |
| Δ*minD2 pTA1392* | 24.26 |
| Δ*minD2* + pSVA3920 | 60.81 |
| Δ*minD2* + pSVA3926 | 23.34 |
| Δ*minD2* + pSVA6011 | 91.96 |
| Δ*minD2* + pSVA6010 | 60 |
| Δ*minD2* + pSVA6037 | 77 |
| Δ*minD2* + pSVA6059 | 21 |
| Δ*hvo_0596* + pTA1392 | 96.5 |
| Δ*hvo_0596minD2* + pTA1392 | 23.79 |
| Δ*CetZ5* +pTA1392 | 99 |
| Δ*CetZ5minD2* +pTA1392 | 24,55 |
| Δ*CetZ6* +pTA1392 | 99 |
| Δ*CetZ6minD2* +pTA1392 | 35* |

Table 5: syntTax report of conserved MinD2 and HVO_0596 proteins

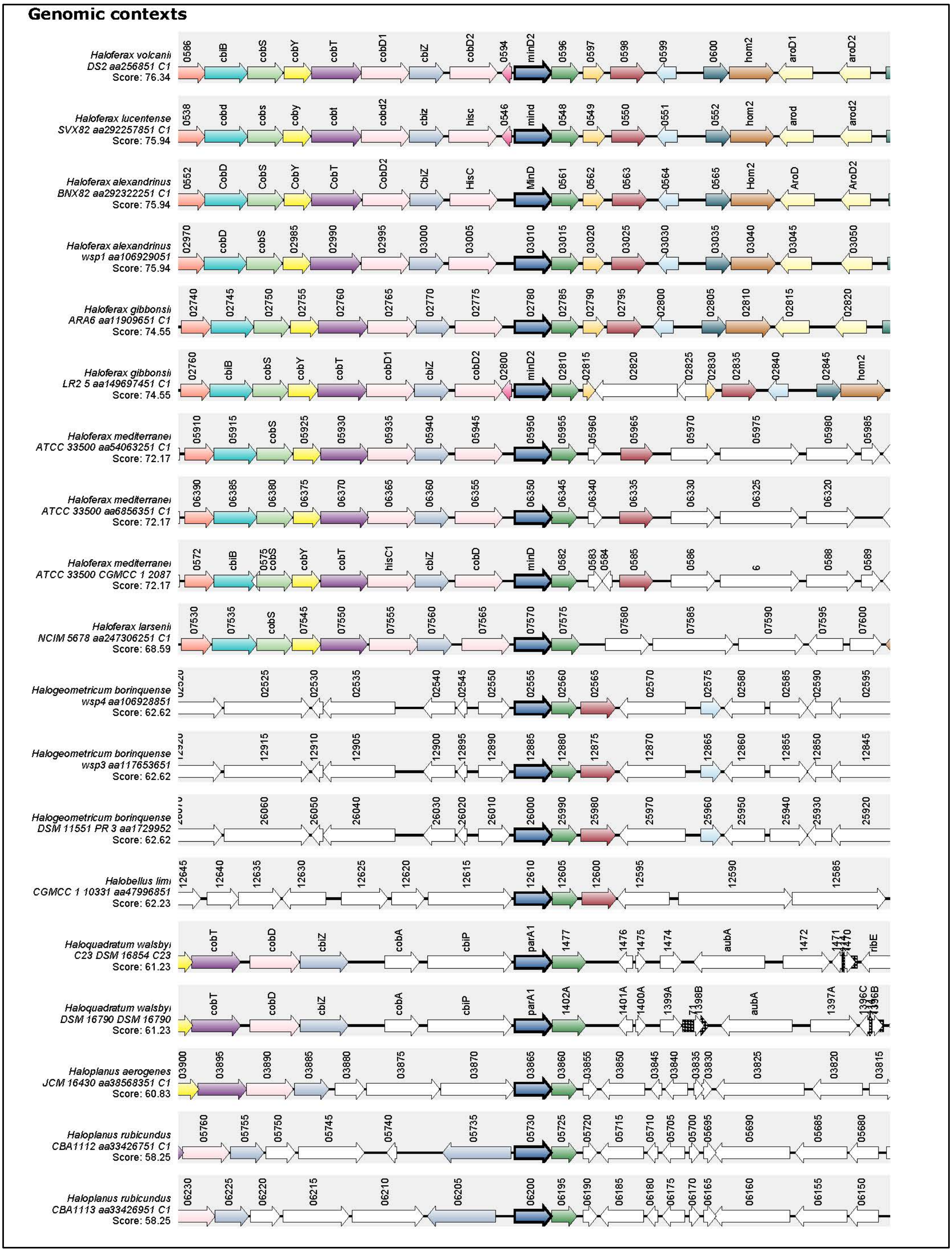
